# ABO blood group, glycosyltransferase activity and risk of Venous Thrombosis

**DOI:** 10.1101/2020.03.01.971754

**Authors:** Manal Ibrahim Kosta, Pascal Bailly, Monique Silvy, Noemie Saut, Pierre Suchon, Pierre-Emmanuel Morange, Jacques Chiaroni, David-Alexandre Trégouët, Louisa Goumidi

## Abstract

**Introduction:** ABO blood group influence the risk of venous thrombosis (VT) by modifying A and B glycosyltransferases (AGT and BGT) activities that further modulates Factor VIII (FVIII) and von Willebrand Factor (VWF) plasma levels. The aim of this work was to evaluate the association of plasma GTs activities with VWF/FVIII plasma levels and VT risk in a case-control study.

**Materials and Methods:** 420 cases were matched with 420 controls for age and ABO blood group. GT activities in plasma were measured using the quantitative transfer of tritiated N-acetylgalactosamine or galactose to the 2’-fucosyl-lactose and expressed in disintegration per minute/30µL of plasma and 2 hours of reaction (dpm/30µL/2H). FVIII and VWF plasma levels were respectively measured using human FVIII-deficient plasma in a 1-stage factor assay and STA LIATEST VWF (Diagnostica Stago).

**Results:** A and B GT activities were significantly lower in cases than in controls (8119±4027 vs 9682±4177 dpm/30µL/2H, p=2.03 × 10^−5^, and 4931±2305 vs 5524±2096 dpm/30µL/2H, p=0.043 respectively). This association was observed whatever the ABO blood groups. The ABO A1 blood group was found to explain∼80% of AGT activity. After adjusting for ABO blood groups, AGT activity was not correlated to VWF/FVIII plasma levels. Conversely, there was a moderate correlation (ρ∼0.30) between BGT activity and VWF/ FVIII plasma levels in B blood group carriers.

**Conclusion:** This work showed, for the first time, that GT activities were decreased in VT patients in comparison to controls with the same ABO blood group. The biological mechanisms responsible for this association remained to be determined.

## Introduction

Venous thrombosis (VT), including deep vein thrombosis (DVT) and pulmonary embolism (PE) is a multifactorial disease whose occurrence and development depend on the interplay of environmental and genetic factors^1^. ABO blood group is one of the first identified genetic risk factors for VT^2^. Recent studies^3–5^ have clarified this association and demonstrated that A1 and B blood groups were at ∼2 fold increased risk of VT compared to O and A2 blood groups.

Despite this convincing association the exact mechanisms relating ABO to VT is not completely characterized. The ABO blood groups are determined by the *ABO* locus located on chromosome 9 (9q34.1-q34.2). The *ABO* locus codes for two glycosyltransferases (GT), A and B, whose role is to transfer a saccharide unit to polypeptides and membrane glycolipids in the red blood cells^6^. AGT and BGT are able to catalyze the specific transfer of a saccharide in activated form, N-acetylgalactosamine (UDP-GalNAc) or D-galactose (UDP-Gal) respectively, on an accepting substrate, the H antigen of minimal structure L-Fuc1-2Gal, which is thus converted into antigenic structure A or B. AGT and BGT are present in Golgi (membrane form) and plasma (soluble form), and are encoded by different alleles at the *ABO* locus. While AGT is encoded by the rs579459 polymorphism at the origin of the A1 blood group, BGT is encoded by the rs8176743 polymorphism that leads to the B blood group. A2 blood group defined by the rs1053878 is characterized by the deletion of one single nucleotide causing a shift in the reading frame leading to a drastic reduction in AGT activity^7^. By contrast, the *ABO* O form codes for an inactive enzyme resulting of a single nucleotide deletion (rs8176719), with individuals of blood group O expressing only the H antigen^6^.

The most likely hypothesis that is generally put forward to explain the relationship between VT and *ABO* is that, by modifying GT expressions, *ABO* participates in controlling the degree of glycosylation of Willebrand factor (VWF), and consequently in its clearance and/or cleavage by ADAMTS13^8,9^. This is supported by the observation of a strong association between VWF levels and the degree of loading of VWF with A and B antigens^10,11^. This would explain the robust association found between the ABO blood group and circulating concentrations of VWF^12^ and factor VIII (FVIII) of which VWF is the transport protein. Individuals with A1 and B blood groups have on average 20% higher circulating VWF and FVIII levels than those with O or A2^13,14^ groups, high plasma levels of VWF and FVIII being associated with increased VT risk^15–18^.

However, the effect of the *ABO* locus on VT risk cannot be explained only by a VWF dependent mechanism. Indeed, it has been shown that the association between the ABO blood group and VT persists after adjusting for FVIII or VWF levels^19^. Besides, since ABO antigens are also expressed on various other tissues including platelets and the vascular endothelium^20^, it could be speculated that the *ABO* locus may modulate VT risk via VWF independent mechanisms. Such hypothesis is supported by several studies reporting the association of the *ABO* locus with different cardiovascular endophenotypes such as intercellular adhesion molecule-1 (ICAM-1), P-selectin and E-selectin^21–25^. Finally, the observation that plasma AGT activity can vary by a factor 1 to 3 in individuals of the same A1 blood^10^ group strongly suggests that the variability in plasma AGT activity cannot be attributable only to ABO blood groups, emphasizing the need for a better assessment of the role of GTs in relation to VT risk.

This work was designed to investigate the association of plasma GTs activities with VWF/FVIII plasma levels and with VT risk and determine whether such associations are fully mediated by the effect of the ABO blood group.

## Material and Methods

### Study description

We conducted a case-control study where patients with VT were compared to healthy donors for their AGT and BGT plasma activities.

Patients were selected from the MARTHA (MARseille Thrombosis Association study) population, a VT patients population extensively described elsewhere^26,27^ and composed of unrelated Caucasian patients with a personal documented history of VT (DVT and/or EP) in the absence of a major VT risk factor (natural coagulation inhibitor deficiency, homozygosity for factor V Leiden or for factor II G20210A mutation, antiphospholipid syndrome) and who consulted at the Centre d’ Exploration des pathologies Hémorragiques et Thrombotiques (CEHT) at La Timone Hospital in Marseille between January 1994 and October 2008. All study subjects were interviewed by a physician about their medical history, which emphasized manifestations of DVT and PE using a standardized questionnaire. The date of occurrence of every episode of VT and the presence of precipitating factors (such as surgery, trauma, prolonged immobilization, pregnancy or puerperium, and oral contraceptive intake) were collected. VT was classified as provoked when occurring within 3 months after exposure to exogenous risk factors, including surgery, immobilization for 7 days or more, oral contraceptive use, pregnancy and puerperium, trauma of the lower limb, long travel (by car > 10 hours, by plane > 5 hours). In the absence of these risk factors, VT was defined as unprovoked. The thrombotic events were documented by venography, Doppler ultrasound, spiral computed tomographic scanning angiography, and/or ventilation/perfusion lung scan.

For the present study, 420 MARTHA participants with available plasma and sampled between December 1994 and June 2015, were phenotyped for AGT and BGT activities. Description of main clinical characteristics of VT patients are reported in supplementary Table S1.

MARTHA patients were compared with 420 controls that were healthy blood donors recruited between February and December 2015 at the Etablissement Français du Sang Alpes-Méditerranée (EFS-AM) with no personal history of cardiovascular disease including VT. As the main objective of this study was to assess the association of GTA and GTB activities with the risk of VT while controlling for the confounding effect of the ABO blood group, controls were selected to match to cases for age at blood sampling (accepting a difference of +/− 5 years) and ABO blood groups.

Informed written consent was obtained in accordance with the Declaration of Helsinki. The study was approved by the local Ethics Committee at La Timone Hospital (Marseille, France).

### Biological parameters

Blood samples were collected by antecubital venipuncture into Becton-Dickinson vacutainer tubes containing 0.105 mol/L trisodium citrate.

Platelet poor plasma was obtained by centrifugation at 2500g for 10 minutes, aliquoted, frozen and stored at - 80°C. DNA was extracted from the EDTA tube samples.

Plasma FVIII coagulant activity was assayed in an automated coagulometer (STA-R; Diagnostica Stago, France). VWF antigen was measured with a commercially available enzyme-linked immunosorbent assay kit from Diagnostica Stago.

Plasma determination of AGT and BGT activities was performed as recommended by Keshvara et al^28^. Briefly, 30 µL of each plasma was incubated in a final volume of 100 µL containing ATP (5mM), MnCl2 (14mM for AGT and 21mM for BGT), 2-(N-morpholino) ethanesulfonic acid (MES) 50mM pH 6.5 for AGT and pH 7 for BGT, an acceptor 2’-fucosyllactose (105µM), and a tritiated specific oligosaccharide (25µM) in the form of UDP-N-acetylGalactosamine [6-^3^N(N)] (American Radiolabeled Chemicals. Inc.) for the AGT activity and UDP-Galactose [6-^3^H] (American Radiolabeled Chemicals. Inc.) for the BGT activity. The reaction mixture was incubated at 37°C for 2 hours and then the reaction was stopped by adding 400µL of water. The radiolabeled synthetic product was purified by negative selection on an anion exchange resin (AG®1-X8). The radioactivity produced (specific oligosaccharide transfer control) was measured in disintegration per minute (dpm) by a scintillation counter. The radioactivity measured was proportional to the GT activity. The activity measurements were carried out in duplicate. Cases and controls were mixed to avoid any batch effect. O blood group carriers were taken as negative controls and the radioactive signal measured for these samples was considered as a nonspecific substrate degradation. In the manuscript, the GT activities are reported in dpm for 30µl of plasma and 2 hours of reaction and individual values were calculated as the mean over duplicates.

ABO blood group was genetically determined using the light cycler technology (Roche Diagnostics, Indianapolis, IN) as previously described^26^. Briefly, the rs8176719-G/deletion, rs1053878-G/A and rs8176743-C/T alleles were genotyped to define ABO O1, A2 and B blood groups, respectively. If none of these alleles were present in an individual, the latter was considered of ABO A1 blood group.

### Statistical analysis

Continuous variables were described by mean and standard derivation while categorical variables were expressed in counts and percentages. ANOVA test was performed to compare continuous variables while a Pearson χ^2^ test was performed to compare categorical variables. A and B GT activities were not normally distributed in contrast with their residues allowing the use of multiple linear regression models in association analysis. The covariates were age, gender and BMI. The statistical significance was assigned at a value of p=0.05. All statistical analyses were performed using SAS 9.4 software.

## Results

### Characteristics of the cases and controls population

Table 1 presents the main characteristics of the 420 cases (Supplementary Table S1) and their matched controls. According to the design of the study no significant difference was observed in age and ABO blood groups distribution. While there was no difference in BMI between cases and controls cases (25 ± 3.3 vs 25 ± 4.9 kg/m^2^, p= 0.99), there was a significant (p= 0.002) higher proportion of women in cases than in controls (53% vs 42%). Factor V Leiden and factor II G20210A mutations were more frequent in cases than controls (27% vs 5%, p= 6.04 × 10^−18^ and 11% vs 3%, p= 1.27 × 10^−5^ respectively). As expected, FVIII and VWF plasma levels were significantly higher in cases than in controls (FVIII: 138 ± 52 IU/dL vs 115 ± 33 IU/dL, and VWF: 144 ± 65 IU/dL vs 116 ± 35 IU/dL, p= 1.01 × 10^−12^ and p= 4.11 × 10^−14^, respectively). After adjustment for sex and age at sampling, this difference was significant in each ABO blood group except in A1A1 and A2B/A2O for VWF and FVIII and in OO for FVIII (Supplementary Tables S2 and S3). Interestingly, these differences remained significant after adjustment for A or B GT activities suggesting that these differences are independent of A and B GT activities. The median time between VT and sampling was 8.5 months with a range of 0.1 to 553 months. There was no significant correlation between AGT (ρ= 0.12, p= 0.06) nor BGT (ρ= 0.03, p= 0.78) and the sampling time after the VT event after adjusting for age at sampling, sex and BMI. At time of sampling, 166 (40%) cases were on anticoagulant treatment and their AGT and BGT. Differences in AGT and BGT activities remained between controls and cases when patients under anticoagulant treatment were excluded from analysis (p= 1.32 × 10^−9^ and p= 0.0041, respectively).

**Table 1:** Main clinical and biological characteristics of the studied population

|  | **Controls**<br>**N=420** | **Cases**<br>**N=420** | ***p*** |
| --- | --- | --- | --- |
| Females | 177 (42) | 222 (53) | 0.002 |
| Age at sampling (years) | 43 ± 12 | 43 ± 13 | 0.46 |
| BMI (kg/m<sup>2</sup>) | 25 ± 3.3 | 25 ± 4.9 | 0.99 |
| FV Leiden carriers | 20 (5%) | 111 (27%) | 6.04 x 10<sup>-18</sup> |
| FII 20210A carriers | 13 (3%) | 45 (11%) | 1.27 x 10<sup>-5</sup> |
| ABO(H) blood group: |  |  |  |
| A1A1 | 19 (5) | 19 (5) | 1 |
| A1A2 | 6 (1) | 6 (1) |  |
| A1B | 26 (6) | 26 (6) |  |
| A1O | 181 (43) | 181 (43) |  |
| A2B | 6 (1) | 6 (1) |  |
| A2O | 28 (7) | 28 (7) |  |
| BB | 1 | 1 |  |
| BO | 68 (16) | 68 (16) |  |
| OO | 85 (20) | 85 (20) |  |
| FVIII levels (IU/dL) | 115 ± 33 | 138 ± 52 | 1.01 x 10<sup>-12</sup> |
| VWF levels (IU/dL) | 116 ± 35 | 144 ± 65 | 4.11 x 10<sup>-14</sup> |
| AGT dpm/30μL | 9682 ± 4177 | 8119 ± 4027 | 2.03 x 10<sup>-5</sup> |
| BGT dpm/30μL | 5524 ± 2096 | 4931 ± 2305 | 0.043 |
| Anticoagulant treatment |  | 166 (40) |  |
| Time between VT and sampling (months) |  | 8.5 (0.1 – 553) |  |

N: number of individuals; Age at sampling, BMI, FVIII and VWF are reported in mean + SD; Time between VT and sampling is reported in median and range; Qualitative variables are reported in N (%); *p*: adjusted for age at sampling, sex and BMI

Of note there was a moderate but significant (ρ= - 0.13, p= 0.03) negative correlation between BMI and AGT in cases, but not in controls (ρ= - 0.02, p= 0.74). No correlation was observed with BGT (data not shown). Finally, after adjusting for age at sampling and BMI, we did not observe any significant difference in AGT between males and females (8655 ± 4015 vs 9165 ± 4329 dpm/30µL/2H, p= 0.57). The same hold for BGT (5051 ± 2196 in males vs 5400 ± 2235 dpm/30µL/2H in females, p= 0.22).

### Comparison of A and B GT activities between cases and controls

Results of the case-control association analysis of AGT and BGT activities are detailed in Table 2. It is important to remember that AGT can be quantified only in carriers of at least one A1 or A2 allele keeping in mind that the A2 allele is associated with very low AGT compared to A1 allele^7^. Similarly, BGT can only be quantified in carriers of at least one B allele. In OO carriers, GT is not functional and corresponding activities cannot be measured. In individuals heterozygotes for the O allele (i.e A1O, A2O or BO), measured GT results from the GT activity produced by the second allele present (i.e A1, A2 or B).

**Table 2:**
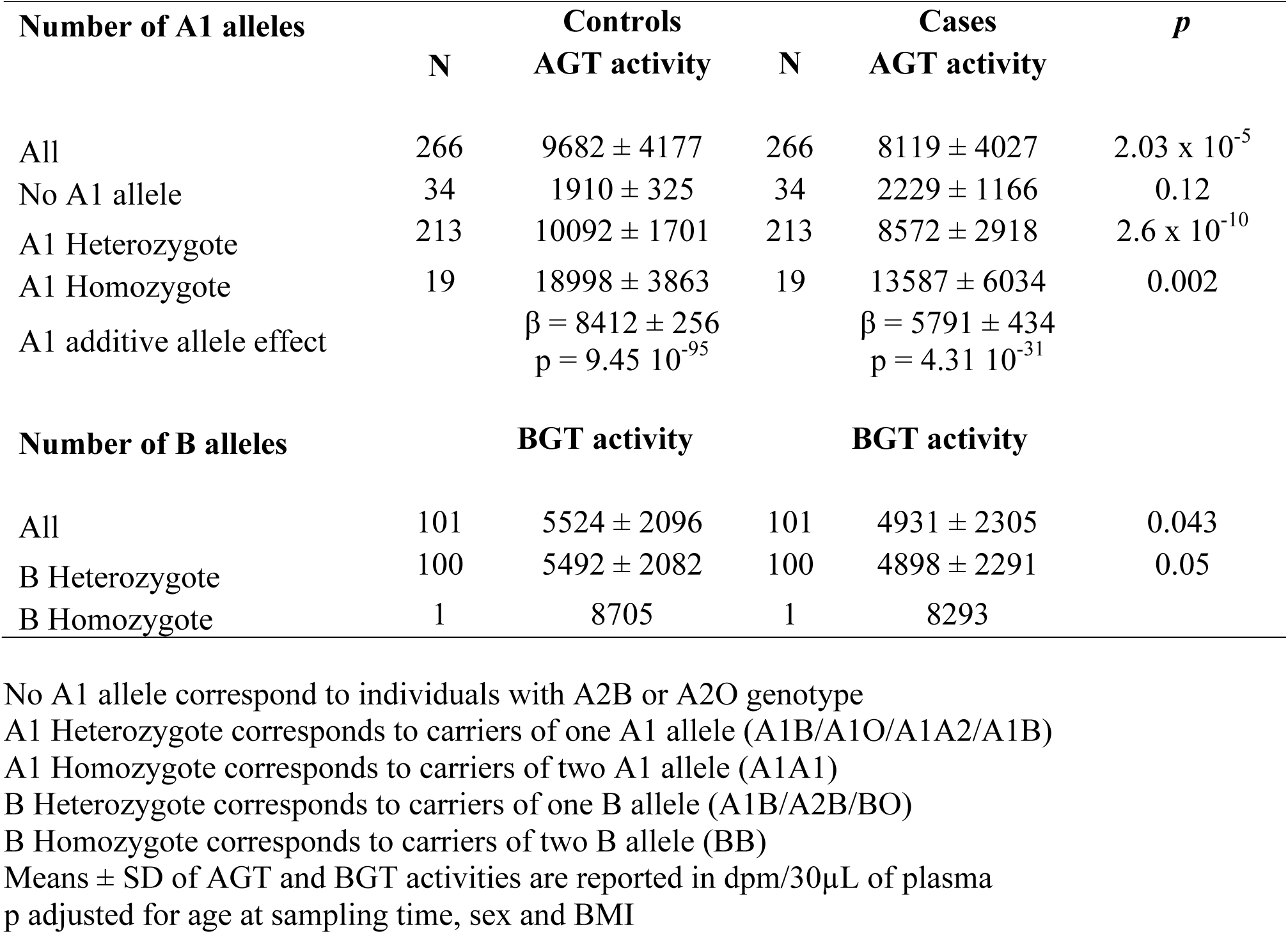
Comparison of A and B GT activities between controls and cases according to ABO blood groups

After adjusting for age at sampling, sex and BMI, A and B GT activities were significantly lower in cases than in controls (8119 ± 4027 vs 9682 ± 4177 dpm/30µL/2H, p= 2.03 × 10^−5^, and 4931 ± 2305 vs 5524 ± 2096 dpm/30µL/2H, p= 0.043, respectively). And these associations were observed whatever the ABO blood group where the corresponding GT could be quantified except for no A1 allele group which reflects the A2 GT activity (p= 0.12, Table 2).

As expected, plasma AGT activity increased with the number of A1 allele, under a fairly additive model, both in controls and in cases (Table 2). The allelic effect of the A1 allele on AGT was 8412 ± 256 (p= 9.45 10^−95^) in controls and 5791 ± 434 (p= 4.31 10^−31^) in cases. In controls, the A1 allele was found to explain about 80% of the plasma AGT variability.

Association of BGT activity with the number of B allele could not be tested because of low number of individuals (n= 2, 1 case and 1 control) of BB genotype. However, these two individuals were at much higher BGT activity than individual heterozygotes for the B allele (Table 2).

### Correlation analysis of A and B GT activities with VWF and FVIII levels

After adjusting on age at blood collection, sex and BMI, we observed significant correlations between AGT activity and VWF in controls (ρ= 0.24, p= 5 × 10^−5^) and cases (ρ= 0.23, p= 0.0002) and FVIII levels in controls (ρ= 0.22, p= 0.0004) and cases (ρ= 0.25, p= 0.0002). However, in subjects with the same ABO blood group, no correlation between AGT activity and VWF nor FVIII levels is observed as shown in Table 3. This indicates that after adjusting for ABO blood groups, there is no association between AGT activity and plasma levels of VWF and FVIII.

**Table 3:** Correlations between plasma GT activities and plasma levels of VWF and FVIII.

|  |  | VWF |  | FVIII |  |
| --- | --- | --- | --- | --- | --- |
|  |  | Controls | Cases | Controls | Cases |
| **GTA** | No A1 | 0.24 (n= 34)<br>p= 0.18 | 0.46 (n= 34)<br>p= 0.01 | 0.22 (n= 34)<br>p= 0.23 | 0.60 (n= 29)<br>p= 0.002 |
|  | A1 Heterozygotes | 0.09 (n= 212)<br>p= 0.18 | 0.11 (n= 211)<br>p= 0.09 | 0.02 (n= 212)<br>p= 0.80 | 0.04 (n= 173)<br>p= 0.63 |
|  | A1 Homozygotes | -0.12 (n= 19)<br>p= 0.65 | 0.23 (n= 19)<br>p= 0.40 | 0.11 (n= 19)<br>p= 0.67 | -0.25 (n= 13)<br>p= 0.50 |
| **GTB** |  |  |  |  |  |
|  | B Heterozygotes | 0.28 (n= 99)<br>p= 0.004 | 0.18 (n= 98)<br>p= 0.08 | 0.29 (n= 100)<br>p= 0.004 | 0.19 (n= 86)<br>p= 0.09 |

No A1 allele correspond to individuals with A2B or A2O genotype
A1 Heterozygote corresponds to carriers of one A1 allele (A1B/A1O/A1A2/A1B)
A1 Homozygote corresponds to carriers of two A1 allele (A1A1)
B Heterozygote corresponds to carriers of one B allele (A1B/A2B/BO)
B Heterozygotes without A1 corresponds to carriers of one B allele without A1 allele (A2B/BO)
B Heterozygotes with one A1 corresponds to carriers of one B allele and one A1 allele (A1B)
p adjusted for age at sampling time, sex and BMI

In B heterozygotes carriers, after adjusting on age at blood collection, sex and BMI, BGT activity was correlated with VWF levels (ρ= 0.28, p= 0.004) in controls and in cases (ρ= 0.18, p= 0.08). The same pattern hold with FVIII levels (ρ= 0.29, p= 0.004 and ρ= 0.19, p= 0.09) in controls and in cases, respectively.

## Discussion

We here report the first investigation of A and B GT plasma activities in a case-control study for VT. We observed that individuals with a personal history of VT exhibit decreased A and B GT activities than healthy individuals and that these differences were independent of the ABO blood groups. We also observed some variability in AGT/BGT levels among individuals from the same ABO blood group. All these findings strongly suggest the existence of other mechanisms beyond the ABO blood group that determine the plasma GT variability.

Current biological knowledge does not explain the decrease in GT activities observed in VT patients. All biomolecules regulated by GT expression have not yet been fully characterized^11^. Beyond associations with levels of circulating VWF, ABO phenotype is also thought to affect endothelial leucocytes interaction and leucocyte recruitment and transmigration to inflamed endothelium by influencing serum levels of endothelial-derived adhesion molecules, including soluble P-selectin, E-selectin and ICAM-1^21,22,29^. Notably, the direction of ABO blood group association with blood levels of these endothelial-derived adhesion molecules is opposite to that for VWF. This observation underlines the complexity between ABO locus, GT activities and thrombosis. We can thus hypothesize that the higher GT activities in the control cases might have an effect on the shedding, clearance or secretion of molecule(s), yet to be characterized, which protects from VT and thus impairing its activity.

We also clarified the relationship between AGT activity, VWF and FVIII levels and ABO blood groups^11^. O’Donnell et al showed, in healthy volunteers a direct relationship between the number of A allele carried, AGT activity, VWF and FVIII plasma levels^10^. In the present study, we confirmed the dose effect of the A1 blood group on AGT activity. We also observed a dose effect of the A1 blood group on VWF and FVIII levels (Supplementary Tables 2 and 3). However, we demonstrated that after adjusting for ABO blood group, there was no correlation between AGT activities and VWF/FVIII plasma levels. Interestingly, BGT activity was found to correlate with VWF/FVIII plasma levels suggesting the involvement of other factors than ABO blood group in the relationship between BGT and VWF/FVIII plasma levels.

There are several limitations of the present work. First, our study is a case-control study in which the blood sample was collected after the VT event. Therefore, we cannot exclude the possibility that differences in plasma levels of FVIII, VWF and GT activities resulted from the thrombotic event itself. However, the blood draw was performed with a median of 8.5 months after the thrombotic event and no correlation was observed between VWF and the time between the VT event and the blood draw (ρ= −0.04, p=0.43) nor FVIII (ρ= 0.06, p= 0.3). There was also no significant correlation between AGT (ρ= 0.12, p= 0.06) nor BGT (ρ= 0.03, p= 0.78) and the time between the VT event and the blood draw. Moreover, after excluding patients sampled within the 3 months after the VT, the differences remained significant between controls and cases for VWF (117 ± 36 vs 138 ± 59 IU/dL, p= 1.17 × 10^−8^) and FVIII levels (115 ± 33 vs 135 ± 49 IU/dL, p= 3.26 × 10^−8^). Second, plasma from cases and controls were not taken at the same time. The mean duration of storage at −80°C for the cases was 128 months (range 12 to 260 months) and 12 months (range 5 to 21 months) for the controls. Even if the same pre-analytical and frozen conditions were applied to cases and controls samples we cannot exclude an effect of the storage duration on GT activities. However, we found no correlation between the storage duration of the samples and AGT activity in cases carrying no A1 allele, one A1 allele or two A1 alleles (p= 0.07; p= 0.29 and p= 0.66 respectively). Conversely, we found a significant correlation between BGT activity and the storage duration of the samples in cases (ρ= −0.29, p= 0.003). Third, anticoagulant therapy may have influenced the levels of GT activity in cases as 40% of them were on treatment at time of blood sampling. However, after excluding these patients, the case-control difference in GT activity remained significant (p= 1.32 × 10^−9^ for AGT and p= 0.0041 for BGT after full adjustment). Fourth, the specific design of the present study with enrichment of the control group in non-O blood group renders difficult to extrapolate the results to the general population. Last, ABO blood group genotypes were identified in MARTHA patients by genotyping 3 single nucleotide polymorphisms (rs8176719-G/deletion, rs1053878-G/A and rs8176743-C/T to define ABO O1, A2 and B blood groups, respectively).If none of these alleles were present in an individual, the latter was considered of ABO A1 blood group. Although this method allows a good determination of the genotypic blood group in the majority of cases, it does not allow us to rule out genotyping errors.

In conclusion, we reported, for the first time, lower GT activities in patients with VT than healthy controls, independently of the ABO blood group. Further studies, and notably in an unselected population, are needed to confirm these findings and to better characterize the mechanisms underlying these associations.

## Acknowledgments

we thank Jennifer Garcia for technical assistance

