## Supplementary material for "ABO blood group, glycosyltransferase activity and risk of Venous Thrombosis": Supplemnatl_tables

**Table S1: Main characteristics of VT events in the 420 cases**

| | |
|---|---|
| Age at first VT (years) | 37 ± 13 |
| Type of VT: | |
| DVT | 324 (77) |
| PE alone | 41 (10) |
| DVT+PE | 55 (13) |
| Circumstances | |
| Provoked | 279 (65) |
| Surgery | 64 (15) |
| Immobilization | 53 (12) |
| Pregnancy/post-partum | 47 (11) |
| Oral contraceptives/women | 77 (35) |
| Others | 32 (8) |
| Unprovoked | 147 (35) |
| Recurrent VT events | 142 (34) |

N: number of individuals; Quantitative variables are reported in mean ± SD and qualitative variables in N (%)

Surgery in the 3 months before VT event; Immobilization > 7 days; Others: trauma of the lower limb, long travel (by car > 10 hours, by plane > 5 hours), VT on catheter

**Table S2: Factor VIII plasma levels (in IU/dL) in cases and controls according to ABO blood group**

| | Controls | | Cases | | $p^1$ | $p^2$ |
|---|---|---|---|---|---|---|
| | N | mean ± SD | N | mean ± SD | | |
| No A1 | 34 | 102 ± 32 | 39 | 128 ± 42 | 0.008 | 0.04 |
| A1 Heterozygotes | 212 | 118 ± 32 | 173 | 142 ± 49 | $7.76 \times 10^{-8}$ | $1.48 \times 10^{-9}$ |
| A1 Homozygotes | 19 | 130 ± 45 | 13 | 150 ± 68 | 0.58 | 0.55 |
| B carriers | 101 | 122 ± 30 | 87 | 161 ± 52 | $2.50 \times 10^{-9}$ | $1.62 \times 10^{-10}$ |
| OO | 85 | 102 ± 29 | 68 | 114 ± 46 | 0.06 | |

No A1 allele includes individuals with A2B or A2O genotype
A1 Heterozygotes include carriers of one A1 allele (A1B/A1O/A1A2/A1B)
A1 Homozygotes corresponds to carriers of two A1 allele (A1A1)
B carriers corresponds to carriers of one B allele (A1B/A2B/BO) or two B alleles (BB)
N: number of individuals; $p^1$: adjusted for age at sampling and sex; $p^2$: adjusted for age at sampling, sex and GTs activities

**Table S3: VWF plasma levels (in IU/dL) in cases and controls according to ABO blood group**

| | Controls | | Cases | | $p^1$ | $p^2$ |
|---|---|---|---|---|---|---|
| | N | mean ± SD | N | mean ± SD | | |
| No A1 | 34 | 104 ± 32 | 34 | 120 ± 45 | 0.09 | 0.27 |
| A1 Heterozygote | 212 | 123 ± 33 | 211 | 149 ± 59 | $5.54 \times 10^{-8}$ | $7.44 \times 10^{-9}$ |
| A1 Homozygote | 19 | 134 ± 27 | 19 | 163 ± 78 | 0.1 | 0.06 |
| B carriers | 100 | 126 ± 30 | 99 | 167 ± 71 | $2.47 \times 10^{-7}$ | $2.88 \times 10^{-8}$ |
| OO | 85 | 91 ± 34 | 83 | 119 ± 64 | 0.0008 | |

No A1 allele includes individuals with A2B or A2O genotype
A1 Heterozygotes include carriers of one A1 allele (A1B/A1O/A1A2/A1B)
A1 Homozygotes corresponds to carriers of two A1 allele (A1A1)
B carriers corresponds to carriers of one B allele (A1B/A2B/BO) or two B alleles (BB)
N: number of individuals; $p^1$: adjusted for age at sampling time and sex; $p^2$: adjusted for age at sampling, sex and GTs activities
